# Germ Granules Functions are Memorized by Transgenerationally Inherited Small RNAs

**DOI:** 10.1101/576934

**Authors:** Itamar Lev, Itai Antoine Toker, Yael Mor, Anat Nitzan, Guy Weintraub, Ornit Bhonkar, Itay Ben Shushan, Uri Seroussi, Julie M. Claycomb, Hila Gingold, Ronen Zaidel-Bar, Oded Rechavi

**Affiliations:** Department of Neurobiology, Wise Faculty of Life Sciences & Sagol School of Neuroscience, Tel Aviv University, Tel Aviv, Israel 69978; Department of Cell and Developmental Biology, Sackler School of Medicine, Tel Aviv University, Tel Aviv, Israel 69978; Deptartment of Molecular Genetics, University of Toronto

## Abstract

In *C. elegans* nematodes, components of liquid-like germ granules were shown to be required for transgenerational small RNA inheritance. Surprisingly, we show here that mutants with defective germ granules (*pptr-1*, *meg-3/4*, *pgl-1*) can nevertheless inherit potent small RNA-based silencing responses, but some of the mutants lose this ability after many generations of homozygosity. Animals mutated in *pptr-1*, which is required for stabilization of P granules in the early embryo, display extremely strong heritable RNAi responses, which last for tens of generations, long after the responses in wild type animals peter out. The phenotype of mutants defective in the core germ granules proteins MEG-3 and MEG-4, depends on the genotype of the ancestors: Mutants that derive from maternal lineages that had functional MEG-3 and MEG-4 proteins exhibit enhanced RNAi inheritance for multiple generations. While functional ancestral *meg-3/4* alleles correct, and even potentiates the ability of mutant descendants to inherit RNAi, defects in germ granules functions can be memorized as well; Wild type descendants that derive from lineages of mutants show impaired RNAi inheritance for many (>16) generations, although their germ granules are intact. Importantly, while P granules are maternally deposited, wild type progeny derived from *meg-3/4* male mutants also show reduced RNAi inheritance. Unlike germ granules, small RNAs are inherited also from the sperm. Moreover, we find that the transgenerational effects that depend on the ancestral germ granules require the argonaute protein HRDE-1, which carries heritable small RNAs in the germline. Indeed, small RNA sequencing reveals imbalanced levels of many endogenous small RNAs in germ granules mutants. Strikingly, we find that *hrde-1;meg-3/4* triple mutants inherit RNAi, although *hrde-1* was previously thought to be essential for heritable silencing. We propose that germ granules sort and shape the RNA pool, and that small RNA inheritance memorizes this activity for multiple generations.

## Introduction

*C.elegans* worms synthesize small RNAs that can regulate mRNA expression and exert physiological changes across multiple generations ^1,2^. Environmental challenges can shape the pool of heritable germline endogenous small interfering RNAs (endo-siRNAs), leaving a mark that persists in the progeny ^3,4^. Similarly, exogeneous administration of double strand RNA (dsRNA) triggers exogenous siRNA-mediated transgenerational silencing of cognate genes ^5^. Typically, dsRNA-triggered heritable responses persist for 3-5 generations ^6,7^, owing to a dedicated pathway which regulates the duration of each heritable response ^7,8^. Maintenance of RNAi interference (RNAi) over generations involves specialized argonaute proteins ^9–11^, amplification of the small RNAs by RNA-dependent RNA Polymerases ^12,13^ (RdRPs), and chromatin modifiers ^8,14–16^. Many of the factors which participate in transgenerational small RNA inheritance reside in germline-specific ribonucleoprotein complexes termed germ granules ^17–23^.

P granules are RNA-rich, cytoplasmic, perinuclear liquid-like condensates ^24^, which are maternally deposited to the oocyte ^25,26^, and following fertilization are asymmetrically segregated to the P lineage designated to become the germline ^27^. P granules are required for maintaining the identify of germ cells throughout development, by preventing ectopic expression of somatic genes ^28–30^. P granules are localized adjacent to nuclear pores, to survey the expression of mRNA molecules exiting the nucleus ^22,31,32^. Various P granules-associated proteins were found to be required for RNAi or for RNAi inheritance, including the P granule core-forming proteins PGL-1 ^33,34^, MEG-3 and MEG-4 ^20^, and the VASA homolog GLH-1 ^35^. Several argonaute proteins are also localized to P granules including PRG-1 ^11^, WAGO-1^19^, CSR-1^17^.

Recently, P granules were found to be part of a tri-condensate assemblage which includes also the Mutator granule (or M granule), and a newly discovered Z granule ^20^. The Z granules co-localize with P granules, but during the mitotic stage in the adult germline the two granules separate, while remaining adjacent to each other ^20^. Z granules contain the WAGO-4 argonaute ^20,21^, and the ZNFX-1 protein ^20,22^, needed for balanced small RNA amplification^22^. Both proteins were shown to be required for RNAi inheritance ^20–22^.

The formation and stabilization of germ granules is a tightly regulated process ^24^. PPTR-1 is a regulatory subunit of the conserved PP2A phosphatase holoenzyme ^36^ that stabilizes maternally deposited P granules in the early embryo, regulating the correct segregation of the granules to the germline blastomere across cell divisions ^37^. In *pptr-1* mutants, P granules are evenly distributed throughout the embryo, and are reduced drastically in number and size ^37,38^. PPTR-1 acts on MEG-3 and MEG-4, which are intrinsically-disordered serine-rich proteins that function redundantly to stabilize the condensed phase of P granules via binding of RNA and PGL-1 ^39,40^. In comparison to wild type animals, *meg-3/4* zygotes display a ~90% loss of P granules ^39^.

In this manuscript, we examined the contribution of P and Z granules to RNAi inheritance. Surprisingly, we found that the germ granules-defective mutants *pptr-1* and *meg-3/4* exhibit enhanced RNAi inheritance. Our experiments reveal that *pptr-1* mutants consistently display enhanced RNAi inheritance, while the RNAi inheritance capacity of *meg-3/4* mutants is affected by history, as it is determined by the ancestral, rather than the current, *meg-3/4* genotype. While P granules are known to be maternally contributed, we report that germ granules morphological changes are not inherited transgenerationally and therefore these entities do not carry the heritable memory. Instead, we propose that germ granules generate transgenerational changes via shaping of the heritable pool of small RNAs.

## Results

To study the relationship between small RNA inheritance and germ granules, we crossed different germ granule mutants to worms which contain a single-copy *gfp* driven by *Pmex-5*-driven in the germline. This *gfp* transgene is commonly used as a target for RNAi inheritance experiments ^5,7,14,35^. Surprisingly, when we tested *pptr-1* and *meg-3/4* mutants, we found that both strains exhibit enhanced RNAi inheritance. *pptr-1* mutants displayed extremely strong heritable responses, persisting for more than 70 generations (Figure 1A, B and C). Stronger RNAi inheritance was observed also when we used the endogenous *oma-1* gene as a target (**Figure S1**). To test this, we crossed the mutants with the redundant and temperature-sensitive dominant lethal *oma-1(zu405)* allele. Heritable silencing of this allele is necessary for survival of the *oma-1* mutant at the restrictive temperatures ^6^. After we validated previous reports showing that *pptr-1* and *meg-3/4* mutants indeed display defective embryonic germ granules (Figure 1D, Figure 3 and **Figure S2**), we conclude that germ granules integrity is not necessary for RNAi inheritance. To strengthen this conclusion we examined the ability of *pgl-1* mutants that we also crossed to the *gfp* transgene to inherit RNAi. *pgl-1* mutants are also defective in P granules assembly ^41^. Although these worms were reported multiple times to be insensitive to germline RNAi ^33,34^. We found that *pgl-1* mutants are competent in RNAi inheritance (**Figure S3,** see more below). We did not observe strengthening of the heritable responses when comparing *pgl-1* to wild type (**Figure S3**). The Z granule argonaute WAGO-4 was shown to be required for inheritance of dsRNA-derived exogenous siRNAs ^20,21^. Our results show that WAGO-4 is indeed required for RNAi inheritance, but only for initiation of the RNAi response, in the parents which are directly exposed to dsRNA. Specifically, *wago-4(−/−)* mutant progeny that derive from RNAi-exposed *wago-4(+/-)* parents, efficiently inherited RNAi for multiple generations (**Figure S4**). This last result show that the consequences of functional WAGO-4 activity in the Z granules can be memorized in further generations even in its absence. In contrast, as was reported by others in the past, the nuclear argonaute HRDE-1 ^9–11,14^ was found to be absolutely required for maintenance of the heritable response (**Figure S4**).

**Figure 1.**
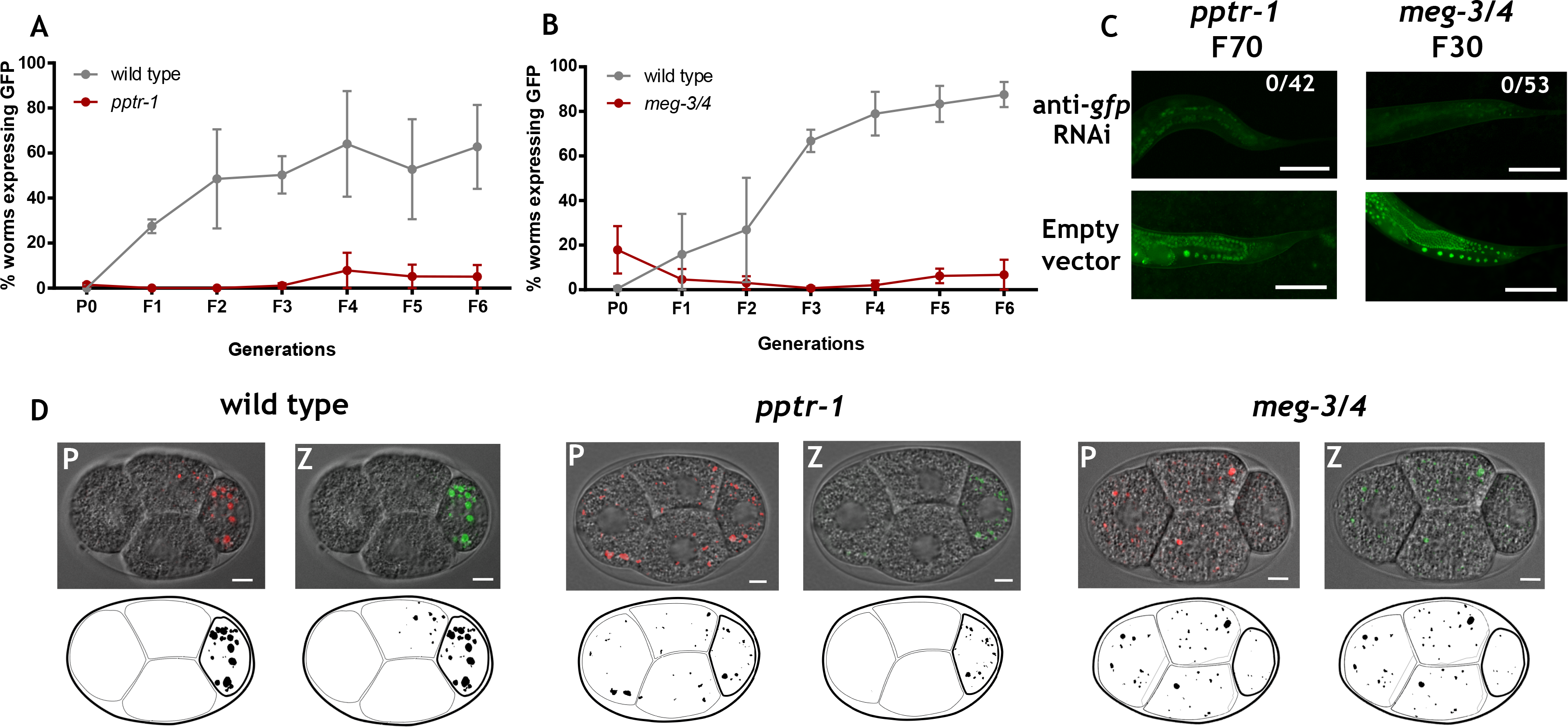
Germ granules mutants exhibit enhanced RNAi inheritance. (**A**) & (**B**) Worms of the indicated genotypes containing a transgene expressing GFP in the germline (*Pmex-5::gfp)* were exposed to dsRNA complementary to *gfp*, to initiate gene silencing via RNAi. The percentage of worms expressing GFP (y-axis) was assessed over generations (x-axis). Shown are mean ± SD from three independent experiments. (**C**) Representative pictures of the germline in the descendants of mutated worms treated with *gfp* RNAi (up) or empty vector. The genotype and generation after RNAi initiation are indicated. Bar: 80 μm. (**D**) Representative pictures of embryos expressing *pgl-1::rfp* (left) and *wago-4::gfp* (right), and their corresponding binarized representations further used for analysis (bottom). The genotypes of the embryos are indicated. Bar: 5 μm.

We noticed that the ability of *meg-3/4* double mutants and *pgl-1* mutants to inherit RNAi was gradually lost after multiple generations, in homozygous mutants that derived from heterozygous parents. Eventually these mutants lost the ability to inherit RNAi, and we reasoned that in previous studies researchers might have assayed mutants that have been homozygous for many generations (**Figure S3**).

To test this hypothesis, we outcrossed *meg-3/4* mutants that were homozygous for multiple generations, and were now refractory to RNAi inheritance. As germ granules are maternally deposited, we performed the cross in two different ways, crossing-in wild type functional *meg-3/4* alleles either via the maternal germline or via the paternal germline (Figure 2A). We isolated F2 *meg-3/4(−/−)* homozygous mutants and F2 *meg-3/4(+/+)* wild type progeny, and examined the capacity of their progeny to initiate RNAi inheritance responses (Figure 2A). As will be elaborated on below, our results clearly demonstrate that the ability of the progeny to inherit RNAi is determined by the genotype of the ancestors, and that the maternal and paternal lineages have different contributions.

**Figure 2.**
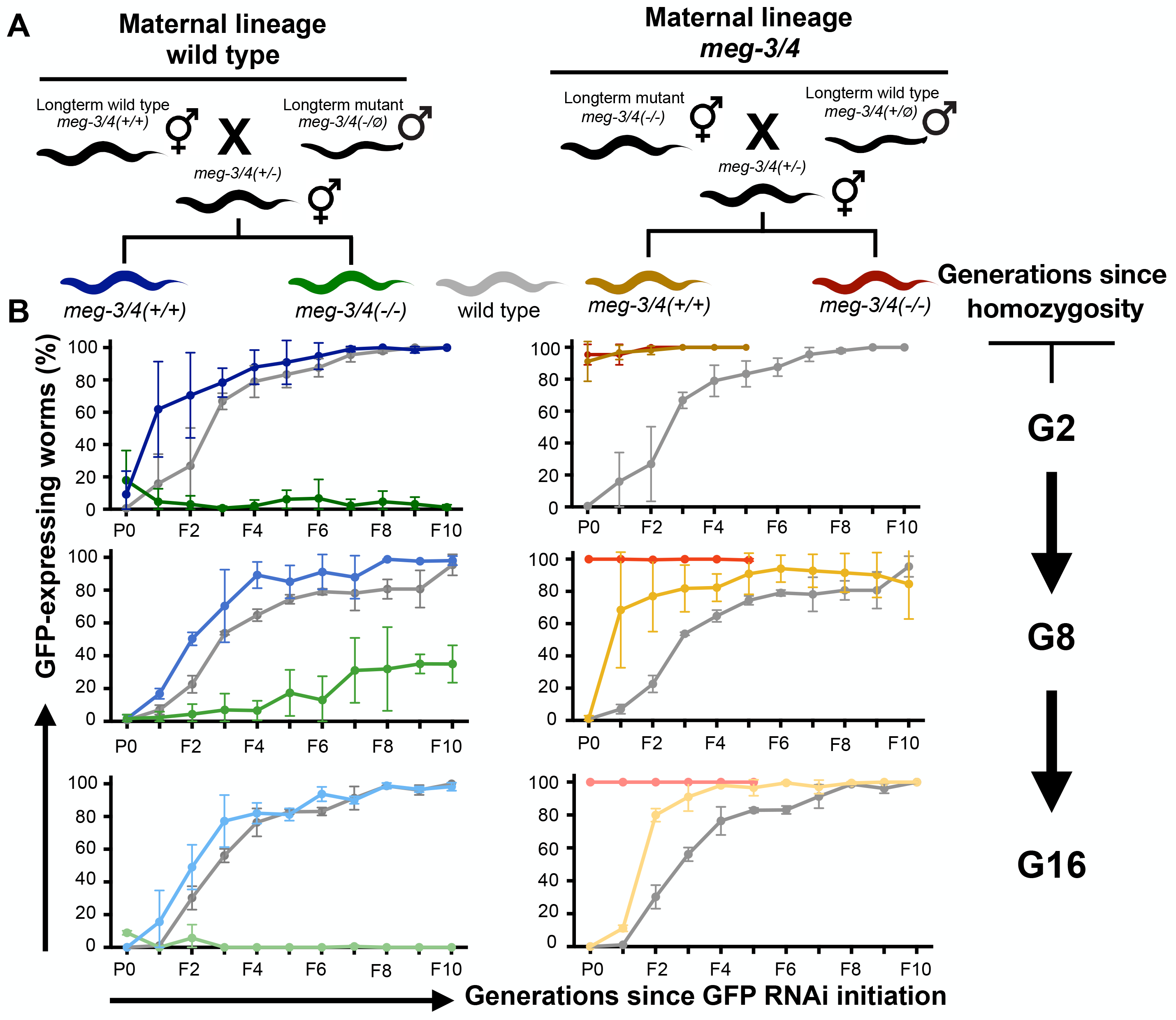
The ancestral germ granules function determines the ability of the progeny to inherit RNAi. **(A)** Schematic diagram depicting the crosses performed to determine the ancestral contribution to RNAi inheritance in the descendants. Long-term *meg-3/4* mutants (hermaphrodites and males) were outcrossed using wild type worms of the opposite sex. All worms contained a transgene expressing GFP in the germline (*Pmex-5::gfp)*. **(B)** Homozygote descendants of the crosses depicted in (A) were exposed to dsRNA complementary to *gfp*, to initiate gene silencing via RNAi. To test the transgenerational dynamics of the ability to response to RNAi, naive descendants were exposed to dsRNA at different time points, both 2 (upper panels), 8 (middle panels) and 16 (lower panels) generations following homozygosity. The proportion of GFP-expressing worms (y-axis, ~50 worms per group) was determined over ten generations (x-axis) following exposure to RNAi. The colors depict the genotypes and lineages according to the scheme in (A). Shown are mean ± SD from two independent experiments (two independent ancestral crosses). In each row (initiation at different generations following homozygosity), all groups were tested side by side, and therefore the same values for the wild type control groups are displayed on both left and right panels. Wild type and *meg-3/4(−/−)* in the maternal wild type lineage for the G2 generation (top left panel) are replicates from figure 1B, for convenience of visualization purposes only.

**Figure 3.**
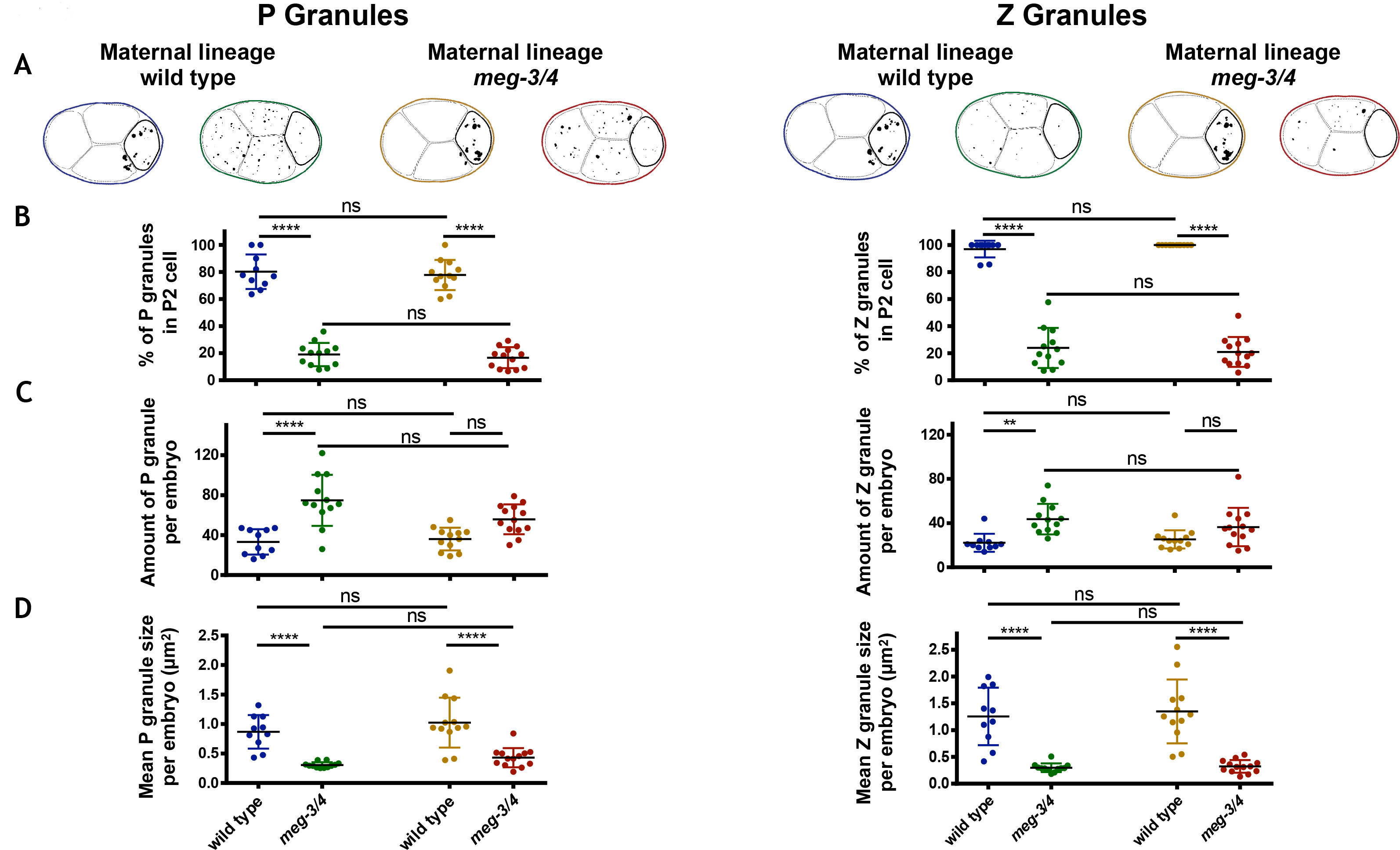
The morphology of the germ granules in the progeny is not affected by the ancestral genotype. Genotypes and maternal lineages of different tested groups are encoded in colors similarly to figure 2. (**A**) Representative binarized pictures of embryos at the 4-cell stage with fluorescent P granules (left) and Z granules (right). The P2 cell (germline lineage) is emphasized with a thicker line. (**B, C, D**) Characterization of germ granules in embryos at the 4-cell stage. Each dot represents one analyzed embryo. Bars represent mean ± SD. P values were determined via Two-way ANOVA with Tukey post-hoc correction for multiple comparisons. ****-p<10^−4^, **-p<0.01, ns-p>0.05. (**B**) Percentage of granules in the P2 cell out of total granules. (**C**) Amount of granules per embryo. (**D**) Mean size of granules per embryo.

When the functional *meg-3/4* alleles were segregated via the maternal lineage, we found that both mutants and wild type grandchildren were competent for RNAi inheritance (the mutant grandchildren even displayed enhanced RNAi inheritance, see Figure 2B, **left panel**). In contrast, when the grandfather contributed the functional *meg-3*/4 alleles, the mutant progeny did not respond to RNAi at all (Figure 2B, **right panel**). Moreover, even the wild type progeny that derive from this cross lost their RNAi inheritance ability, and recovered only after multiple generations (16 generations were not enough for full recovery, Figure 2B, **right panel**). In summary, regardless of their own genotype, the progeny phenocopied the grandmother’s germ granules-controlled RNAi inheritance ability. However, repetitive crossing (a few generations apart) of *meg-3/4* mutants to wild type males did eventually lead to improvement in the capacity of *meg-3/4* to generate RNAi responses (**Figure S5**), suggesting that the male germline can affect this heritable phenotype as well (see more below).

To test if maternal contribution of germ granules could explain the differences described above, we performed similar crosses, and examined if the morphology, abundance and distribution of the germ granules change in the different isolated progeny. In contrast to the RNAi inheritance phenotype of the progeny, which depended on the ancestors’ *meg-3/4* genotype, wild-type progeny had normal germ granules and *meg-3/4* progeny had defective germ granules (Figure 3). Moreover, we found that the P granule characteristics are similar in both RNAi responsive and non-responsive progeny of the same lineage, and the levels of silencing did not correlate with the P granules’ morphology (**Figure S6**). In summary, we could not detect significant differences in the characteristics of the germ granules of F2-F5 isogenic animals that derived from wild type or mutant ancestries.

Germ granules are deposited in the oocyte, not in the sperm. We observed that RNAi inheritance defects can transmit via *meg-3/4(-/Ø)* males to wild type progeny. The defects which transmitted via the paternal lineage lasted for multiple generations, but were weaker in comparison to defects transmitted via the maternal lineage (Figure 2B, **left panel**). Unlike maternally deposited germ granules, small RNAs are inherited from both the sperm and the egg, and therefore we examined whether small RNAs could mediate the transgenerational inheritance of the RNAi defects. The nuclear germline argonaute HRDE-1 is necessary for the inheritance of many endo-siRNAs and dsRNA-derived small RNAs. We found that transient removal of functional HRDE-1 (see scheme Figure 4A) erased the defects inherited from germ granule mutants, and restored the ability of the progeny to inherit RNAi (Figure 4B). Thus, we posit that transgenerational effects which result from defective ancestral germ granules are memorized by inherited small RNAs.

**Figure 4.**
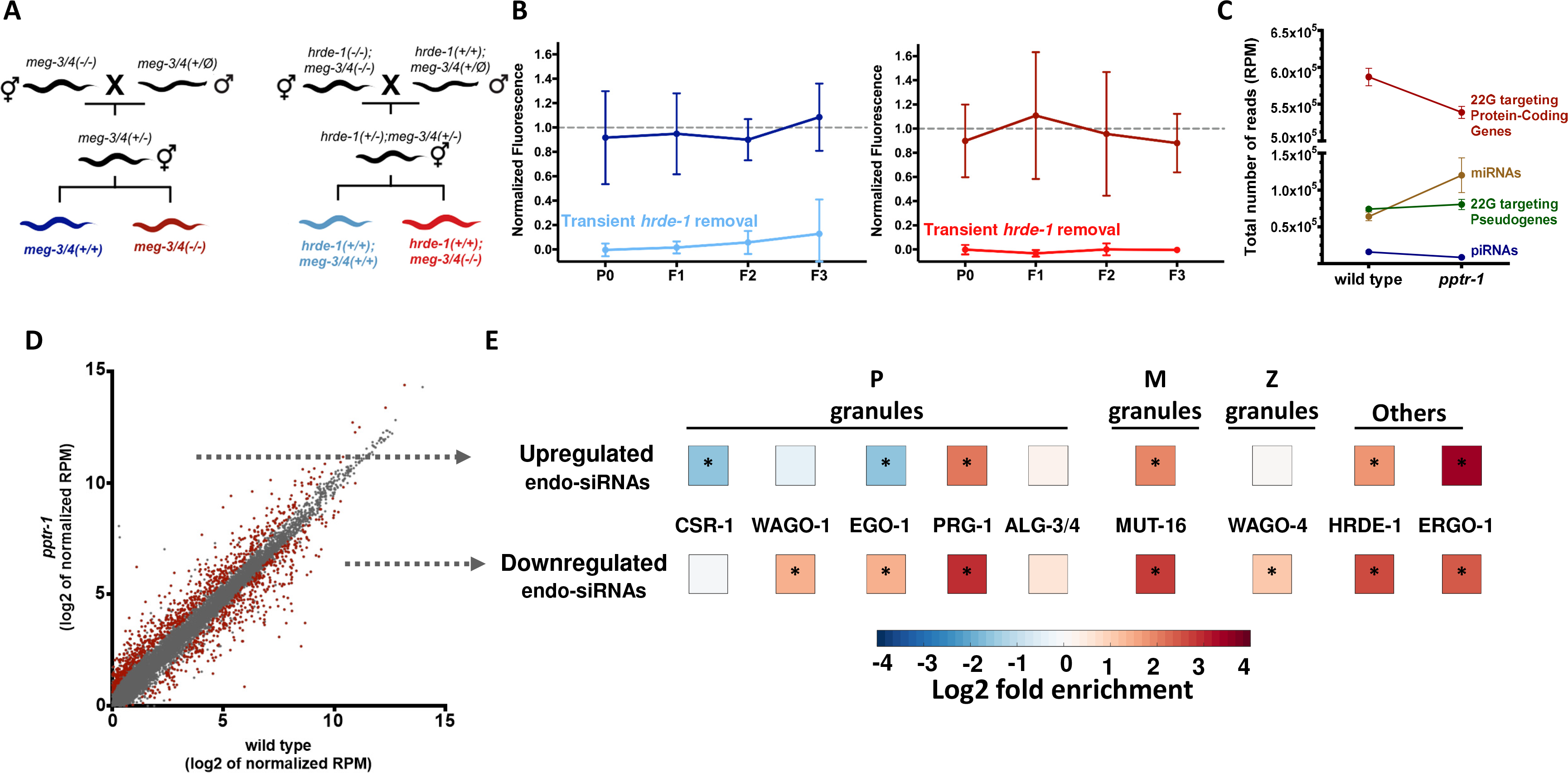
Heritable small RNAs are imbalanced in germ granules mutants and memorize ancestral germ granules defects. (**A**) Schematic diagram depicting the crosses performed to test the ability of transient removal of the *hrde-1* gene, required for inheritance of many small RNAs, to erase defects inherited from germ granule mutants. (**B**) GFP fluorescence (y-axis) of adult worms with the over generations (x-axis), following exposure to *gfp* dsRNA. Genotype and lineage is represented in color according to the scheme in (A). Shown are mean ± SD (30~80 worms per group) normalized to the mean fluorescence value of the corresponding isogenic control worms exposed to empty vector. P values were determined via Two-way ANOVA with Tukey post-hoc correction for multiple comparisons. (**C**) A global analysis of the various small RNA species in dissected germlines of germ granule mutants. Shown are the mean ± SD (3 independent biological repeats) normalized total number of reads (RPM) aligned to the denoted small RNA species in wild type and *pptr-1* mutants. (**D**) A scatter plot analysis of amplified small RNAs. Shown are the mean (log2) normalized reads of small RNAs aligned to each gene in wild type (x axis) and *pptr-1* mutants. Significantly differentially expressed small RNAs (DESeq2, adjusted p-value<0.1) are shown in red. (**E**) An enrichment analysis of differentially expressed small RNAs in germ granules mutants. Shown (color coded) are the enrichment values for small RNAs known to be associated with the denoted small RNA machinery factors. For statistical analysis, the enrichment values of 10,000 random gene sets identical in size to the examined gene set was calculated and ranked (see methods).

Germ granules were hypothesized to balance the levels of different small RNA species ^21,22^, and it is well established that different small RNA pathways compete over limited biogenesis factors ^7,8,42–44^. To understand how germ granules regulate small RNA levels, we sequenced small RNAs from dissected germlines of germ granule-defective *pptr-1* mutants (the mutants that we found to always exhibit strong RNAi inheritance). We found a global reduction in amplified endo-siRNAs targeting protein-coding genes (~22Gs, Figure 4C). Moreover, amplified endo-siRNAs complementary to ~1234 genes were found to be differentially expressed in the mutants (DESeq2 Adjusted p-value<0.1, Figure 4D, **Table S1**). Specifically, we found a disruption in endo-siRNAs that bind the nuclear argonaute HRDE-1 (fold enrichment of 6.2x and 3.4x for downregulated, and upregulated small RNAs in *pptr-1* mutants, p<10^−4^, Figure 4E). Small RNAs that depend on or bind to proteins localized to germ granules were also substantially disturbed, such as endo-siRNAs associated with P granule proteins (the EGO-1 RdRP ^45^, the WAGO-1 ^19^ and the PRG-1 ^46^ argonautes), the M granule protein MUT-16 ^47^, and the Z granule argonaute WAGO-4 ^21^ (Figure 4E).

Our results agree with previous studies suggesting that germ granules are necessary for maintaining small RNA homeostasis and assignment to different argonautes ^21,22^. The HRDE-1 argonaute was shown in numerous studies to be absolutely essential for dsRNA-induced RNAi inheritance ^9–11,14,48,49^. As we found strong RNAi inheritance in germ granule mutants despite severe defects in HRDE-1-associated small RNAs, we hypothesized that when the germ granules are defective, RNAi might be inherited via alternative routes. To test this we performed RNAi inheritance experiments in *hrde-1;meg-3/4* triple mutants. Strikingly, we found that *hrde-1;meg-3/4* triple mutants can inherit RNAi transgenerationally (Figure 5). Together, our results suggest the intriguing possibility that germ granules function in sorting of heritable small RNAs, and that changes in small RNA assortment can persist for multiple generations with extensive consequences to gene regulation in the progeny.

**Figure 5.**
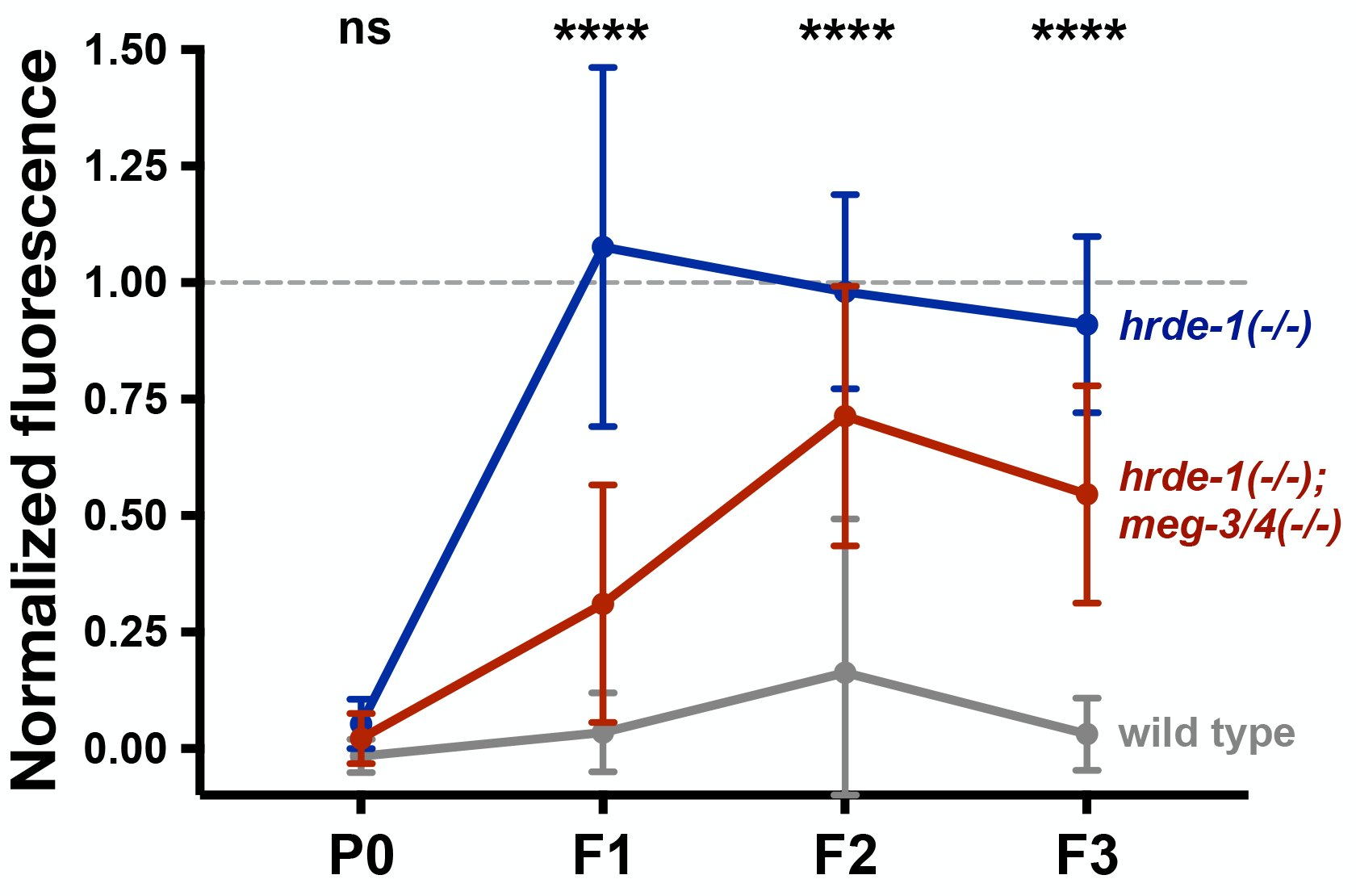
*hrde-1(-/-);meg-3/4(−/−)* triple mutants inherit RNAi transgenerationally. GFP fluorescence (y axis) of adult worms with the denoted genotype over generations (x axis), following exposure to *gfp* dsRNA. Shown are mean ± SD (30~80 worms per group) normalized to the mean fluorescence value of the corresponding isogenic control worms exposed to empty vector. P values were determined via Two-way ANOVA with Tukey post-hoc correction for multiple comparisons. Shown are P values for the comparison of the mean values exhibited by *hrde-1(-/-);meg-3/4(+/+)* worms vs. *hrde-1(-/-);meg-3/4(−/−)* worms. ****-p<10^−4^, ns-p>0.05.

## Discussion

In worms, small RNAs are inherited independently of the DNA sequence ^50^ and enable environmental responses to affect the physiology of multiple generations of progeny. Already more than a 100 years ago it was noted that the cytoplasm is a mixture of liquids in which droplets are suspended ^51^, and RNA-rich granules are present in the germ line of many organisms (more than 80 species were reported to contain such granules, ^52^). Ingenious studies conducted over the years have elucidated the many crucial roles that these granules play, and especially the intriguing ways by which germ granules shape the transcriptome. The realization that germ granules functions change dynamically over the course of multiple generations, could shed new light on germ cell biology, and specifically illuminate the mechanisms that control small RNA-mediated inheritance.

In this manuscript, we revise the current model, demonstrating that mutants defective in germ granules can exhibit potent RNAi inheritance. We hypothesize that the dynamic nature of germ granules activity has not been noted before, since in the past researchers have assayed mutants that have been homozygous for many generations. By performing controlled crosses, we found that the ability of the progeny to inherit RNAi is determined by the genotype of the ancestors, and that the maternal and paternal lineages have different contributions. We found that the heritable effects induced by disruption of germ granules’ functions last for multiple generations (we recorded effects that persist for at least 16 generations in wild type progeny, Figure 2). We suggest a new transgenerational feedback-based model: germ granules shape the pool of heritable small RNAs, and in the next generations small RNAs in turn determine the ability of germ granules to generate *de novo* RNAi inheritance responses.

Several lines of evidence suggest that transgenerational inheritance of small RNAs, rather than multigenerational inheritance of the germ granules themselves, mediates the heritable effects observed following germ granules disruption: (1) the germ granules’ morphology is unaffected by changes to the ancestors’ germ granules, (2) some of the effects can transmit through the paternal lineage, and (3) transient removal of the nuclear argonaute HRDE-1 re-sets the heritable effects of germ granule mutants. Similarly to the interaction described here between heritable small RNAs and germ granules, a transgenerational feed-back exists between small RNAs and other epigenetic pathways, such as chromatin modifications (and DNA methylation in other organisms ^53,54^). These dynamic reciprocal interactions determine eventually how gene-regulatory memories would be inherited across generations ^2,53^.

We suggest that germ granules are not crucial for RNAi *per se*, but rather are important for small RNA homeostasis, with consequences that can persist for multiple generations. Previous studies have shown that the stability of the germ granules is affected by environmental changes (for instance temperature ^55,56^), and also that temperature shifts trigger changes in heritable small RNAs ^3,57^. Because germ granules impact small RNA biogenesis, the granules could be important mediators in the translation of environmental changes to small RNA-mediated transgenerational memory.

More generally, our results show that genetically identical animals can have strikingly different phenotypes, because of their ancestry. First, we provide more evidence that recently outcrossed animals differ from animals that were outcrossed many generations ago due to epigenetic effects. Second, outcrossing using wild type males or hermaphrodites generates different long term phenotypes.

We noticed that while *meg-3/4* mutants eventually become refractory to RNAi inheritance (after many generations), *pptr-1* mutants are consistently responsive to RNAi, and exhibit enhanced RNAi inheritance. The difference between these two mutants can stem from the fact that PPTR-1, a subunit of the PP2A phosphatase holoenzyme, affects additional targets (PP2A has ~460 putative targets in *C. elegans*, ^58^). Additionally, we noticed that the Z granules of *pptr-1* and *meg-3/4* mutants differ. While in *meg-3/4* mutants Z granules fail to segregate towards the P2 cell, these granules segregate normally in *pptr-1* mutants (although the Z granules are smaller in both cases, **Figure S2**).

Finally, prior to Mendel’s realization that traits are transmitted as discreet units of information, it was thought that parents’ heritable materials mix, as liquids do. According to the now obsolete theory known as “Blending Inheritance”, the heritable factors “dilute” in the progeny ^59^. We find that disruption of maternally deposited liquid condensates ^24^ leads to heritable changes which dilute eventually. While the modern mechanistic explanation is obviously different from that used before the re-discovery of Mendel, we propose that the liquid-like properties of RNA-carrying granules lead to deviations from Mendelian inheritance.

## Acknowledgments

We thank all the Rechavi lab members, and especially Sarit Anava, for helpful discussions and scientific guidance. Some strains were provided by the CGC, which is funded by NIH Office of Research Infrastructure Programs (P40 OD010440). We thank Eric Miska and Scott Kennedy for kindly providing us with strains. Special thanks to Dror Cohen for the illustrations that he contributed. O.R is thankful to the Adelis foundation grant #0604916191. This work was funded by ERC grant #335624 and the Israel Science Foundation (grant#1339/17).

## Methods

### Cultivation of worms

All strains were maintained according to standard methods (Stiernagle 2006). Unless noted otherwise, we performed all experiments at 20 degrees. Strains with the oma-1(zu405) temperature-sensitive allele were maintained at 15 degrees, and transferred to 20 degrees (restrictive temperature) for the relevant RNAi experiments. All the strains we used are listed in the key resource table.

### RNAi experiments

HT115 E. coli bacteria bearing the dsRNA-expression bacteria and empty-vector control bacteria were grown overnight in LB medium supplemented with 0.25 μg/ml of Carbenicillin. Bacteria were seeded onto NGM plates supplemented with isopropyl β-D-1-thiogalactopyranoside (IPTG, 1 mM) and Carbenicillin (0.25 μg/ml). A day later, ~10 adult hermaphrodites were placed on the plates, and were allowed to lay eggs for ~4 hours. The progeny laid on this occasion was considered the P0 treated generation. For i*gfp* RNAi inheritance experiments, adult P0 young worms (40~120 per group) were collected for imaging, and for production of the next generation via synchronized egg-laying. For all generations (other than the P0 treated generation), synchronization was performed on the same day of imaging (4 days after egg-laying), by allowing 20 adult worms of each tested group (x2 plates) to lay eggs for ~2 hours on NGM plates seeded with standard OP50 bacteria. For anti-*oma-1* RNAi inheritance experiments, in each generation twelve individual larval (L4) staged worms were placed in individual wells of a 12-well plate. Four days later the number of fertile worms was assessed (at least one progeny) and twelve individual L4 progeny worms were chosen from the most fertile well to continue to the next generation.

Images of the worms were collected using a BX63 Olympus microscope with x10 magnification, and were used to evaluate GFP expression.

### Granules imaging and analysis

Embryos were extracted from young (day 1) adult hermaphrodites. Embryos at the 2-cell stage were chosen and immediately mounted on freshly made 3% agarose pads for imaging. All imaging was carried at 20°C on a Nikon Ti-2 eclipse microscope equipped with a 100X CFI Plan-Apo 1.45 NA objective (Nikon, Tokyo, Japan) and CSU-W1 spinning-disk confocal head (Yokogawa Corporation, Tokyo, Japan). Embryo samples were excited using 488 and 561 nm DPSS-Laser (Gataca, France) and images acquired using Prime95B sCMOS camera (Photometrics, Tucson, AZ). Image acquisition was controlled by MetaMorph software (Molecular Devices, Sunnyvale, CA). We photographed 4 cell stage embryos, using ~30 z stacks per embryo with three channels: Bright field, 488 and 561.

Analysis of granules was carried out using the imageJ2 open source software. First we split the channels and then merged the z stakes using the “Maximal intensity z projection” function. Next, we binarized the fluorescent signal using automated threshold using the “Maximal entropy” method. In the next step we used the “Analyze particles” function to automatically locate the granule foci. Finally, we measured the number and size of the located granule foci. To calculate the percentage of granules in the P2 cell, we manually defined the P2 and embryo regions of interest. Next, we re-applied the “Analyze particles” function to count number of foci in the P2 cell and the whole embryo regions. Foci < 0.1 micron^2^ were excluded from the analysis of P granules.

### Small RNA sequencing

Wild type and *pptr-1(tm3103)* hermaphrodites were collected on the first day of adulthood, washed 4 times in M9 buffer, and transferred in a microscope slide with egg buffer (1M HEPES, 5M NaCl, 1M MgCl2, 1M CaCl2, 1M KCl and 20% tween-20) containing 2mM levamisole. Worms were cut right under the pharynx with a gauge needle, and the evading gonads separated from the body. Gonads were collected from the slide into an eppendorf on ice prior the addition of 300 μl Trizol (Life Technologies). 60μl of chloroform was added to the samples, which were then transferred to a pre-spun Heavy Phase Lock tube and centrifuged at 16,000g for 12 minutes at 4C. We transferred the aqueous phase to a new Heavy Phase Lock tube, and added 1:1 Phenol:Chloroform:Isoamyl Alcohol, before a centrifugation round at 16,000g for 15 minutes at RT. The aqueous phase was transferred to an eppendorf tube, to which we added 1:1 pure ethanol and 20 μg Glycogen (Thermo Fisher), and let incubate overnight in −20C. Samples were then centrifuged at 16,000g for 30 minutes at 4C, then washed twice by removing the supernatant and adding ice-cold 70% ethanol. After the last wash and removal of ethanol, the samples were air dried for 6 minutes, and resuspended in 10μl RNAse-free water. 150 ng of each sample was treated with RNA 5’ polyphosphatase (Epicentre) to ensure small RNA capture independently of 5’ phosphorylation status. Libraries were prepared using NEBNext Multiplex Small RNA Library Prep Set for Illumina (New England Biolabs) according to the manufacturer’s protocol. We measured the samples concentration via Qubit, and tested their quality on an Agilent 2200 TapeStation. Samples were then pooled together and run on a 4% agarse E-Gel (Life Technologies), and the 140-160 nt length bands were excised and purified using MiniElute Gel Extraction Kit. Samples were tested for their quality and concertation on an Agilent 2200 TapeStation and then pooled together. Pools were separated on a 4% agarose E-Gel (Life Technologies) and the 140–160 nt length bands were excised and purified using MiniElute Gel Extraction kit (Qiagen). Libraries were run on an Illumina NextSeq500 sequencer.

### Analysis of GFP silencing

For photographing of the worms ~60 animals were mounted on a 2% agarose slides and paralyzed in a drop of M9 with 0.05% tetramisole. The worms were photographed with 10x objective using a BX63 Olympus microscope (exposure time of 200 ms and gain of 2). Analysis of the photographed was done using Image2 software.

For binary GFP silencing analysis, we manually scored the photographs for the numbers of worms having or lacking any observable germline GFP signal was calculated.

For quantification of GFP fluorescence levels, we manually defined region of the oocytes and regions of three background regions in the quantified worm of each examined worm. Next we calculated a CTCF value as follows: CTCF = Integrated density values of oocyte X – (area of measured oocyte * mean fluorescence of background regions). We normalized the CTCF value to the average CTCF value obtained from photographs of control animals of the same genotype, generation and age which were fed on control plates.

### Quantification of *oma-1* experiment

Each generation we counted the fraction of worms (out of 12) that had progeny.

### Small-RNA Seq analysis

Illumina fastq output files were assessed for quality with FastQC (Simon Andrews 2010). Cutadapt (Martin 2011) was then used for adaptor removal: “*cutadapt -m 15 --discard-untrimmed -a [adaptor sequence] input.fastq*” Clipped reads were aligned to the ce11 assembly of *C. elegans* genome using Shortstack ^60^: “*--mismatches 0 --readfile input.fq genome_reference.fa”*.

Reads were filtered to keep reads of 20 to 23 in length. Next, we counted reads aligning in an anti-sense orientation to genes based on the corresponding Ensembl .gff file, via the HTseq ^61^: *“-m HTSeq.scripts.count --stranded=reverse --mode=intersection-nonempty input.sam GENES.gff”.* We then used the HTseq output file as an input for DESeq2 ^62^.We defined differentially expressed genes using cutoff of adjusted p-value < 0.1.

### Bioinformatic gene enrichment analysis

The enrichment values denote the ratio between the observed representation of a specific gene set within a defined differentially expressed genes group, to the expected one, i.e., the representation of the examined gene set among all protein-coding genes in *C. elegans*. We performed the analysis on 8 gene sets: (1) 1587 targets of HRDE-1 endo-siRNAs ^9^, (2) 4191 targets of CSR-1 ^17^, (3) MUT-16-dependent endo-siRNAs^47^, (4) WAGO-4 targets ^21^, (5) EGO-1-dependent endo-siRNAs ^45^, (6) WAGO-1 targets ^19^, (7) ERGO-1-dependent endo-siRNAs, (8) ALG-3/4 targets ^18^. The putative PRG-1 targets were defined as genes for which, in at least one transcript, the ratio of the # 22G-RNA reads at piRNA target sites between wild type to prg-1 is at least 2 (linear scale).

Enrichment of a given gene set *i* in *pptr-1*-dependent endo-siRNAs was calculated according to the formula:

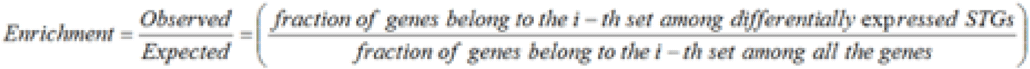

We calculated p values using 10,000 random gene groups identical in size to that of genes targeted by *pptr-1-dependent* endo-siRNAs.

### Bioinformatics analysis of different small RNA species

Briefly, reads were filtered to sizes of 20 -23 in length. Next, reads were counted using HTseq in the sense, and again in the anti-sense orientations based on ce11 Ensembl .gff file. Reads were normalized to the total amounts of aligned reads, and the sum of reads aligning to each small RNA specie was calculated: for small RNAs targeting protein-coding and pseudogenes, reads aligned in the anti-sense orientation were used. For miRNAs and piRNAs, read aligning in the sense orientation were used.

### Statistical analysis

For RNAi experiments and germ granule characterization, statistical tests were performed via GraphPad Prism 6. Details about the tests and corrections for multiple comparisons appear in the corresponding figure legends.

### Data and Software Availability

Raw sequencing files and processed data are available under GEO: GSE128112.

