## Supplemental figures for "Germ Granules Functions are Memorized by Transgenerationally Inherited Small RNAs"

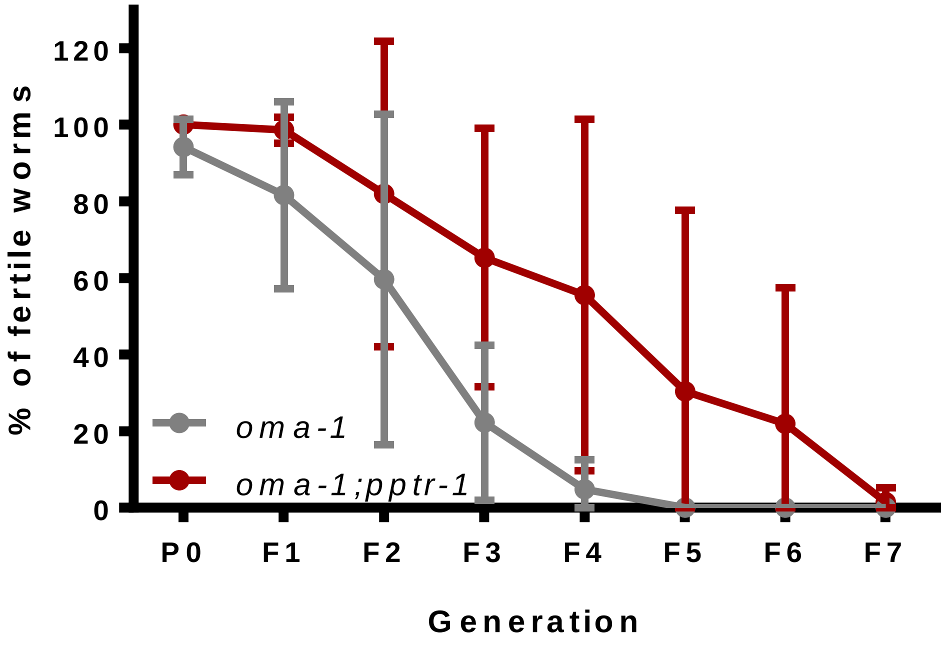


**Figure S1. *pptr-1* mutants exhibit enhanced RNAi inheritance also when silencing the endogenous gene *oma-1***

Transgenerational inheritance of silencing of a temperature-sensitive allele of the endogenous gene *oma-1*. Silencing of *oma-1* in the restrictive temperature is required for fertility. Shown are percentage of fertile worms (y-axis, mean ± SD) from three independent experiments.

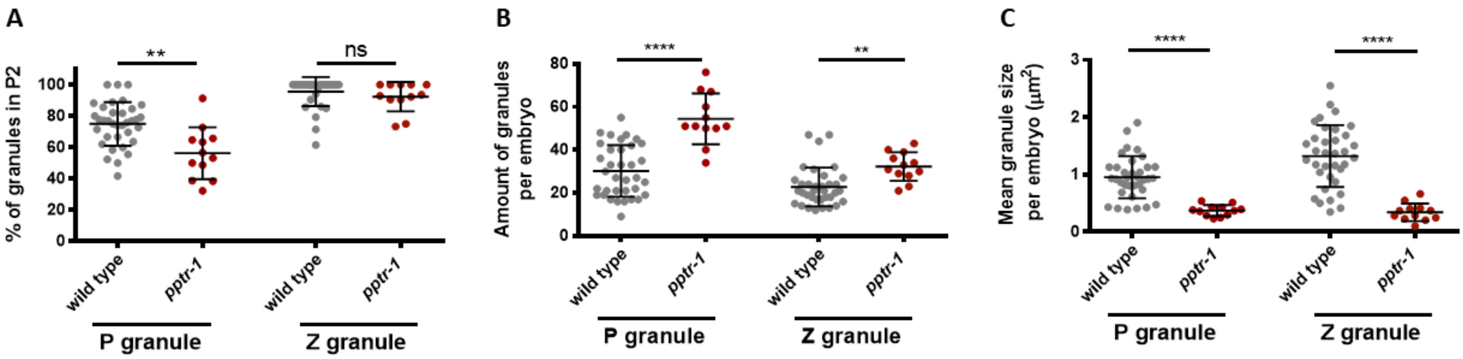


**Figure S2. Analyses of the morphological characteristics of *pptr-1* germ granules.**

Characterization of germ granules in embryos at the 4-cell stage. Each dot represents one analyzed embryo. All available wild type data is displayed, and therefore appears also in Figure 3. Bars represent mean ± SD. P values were determined via Student's two-tailed t-test with Bonferroni post-hoc correction for multiple comparisons. ****- $p<10^{-4}$. **- $p<0.01$. ns- $p>0.05$.

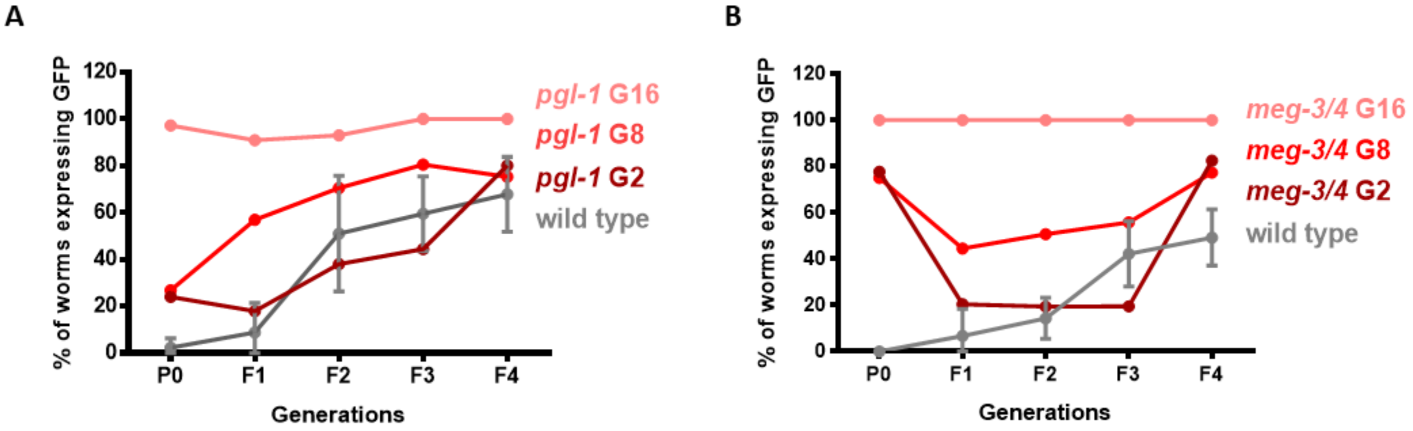


**Figure S3. *pgl-1* and *meg-3/4* can inherit RNAi, but lose this ability after multiple generations.**

Worms of the indicated genotype containing a transgene expressing *gfp* in the germline (*Pmex-5::gfp*) were exposed to *gfp* dsRNA, to initate an RNAi response. The proportion of GFP-expressing worms (y-axis) was measured over generations (x-axis). The tested homozygous mutant strains descend from heterozygotes parents, and “G#” indicates the number of generations that have passed since homozygosity at the time of RNAi initiation.

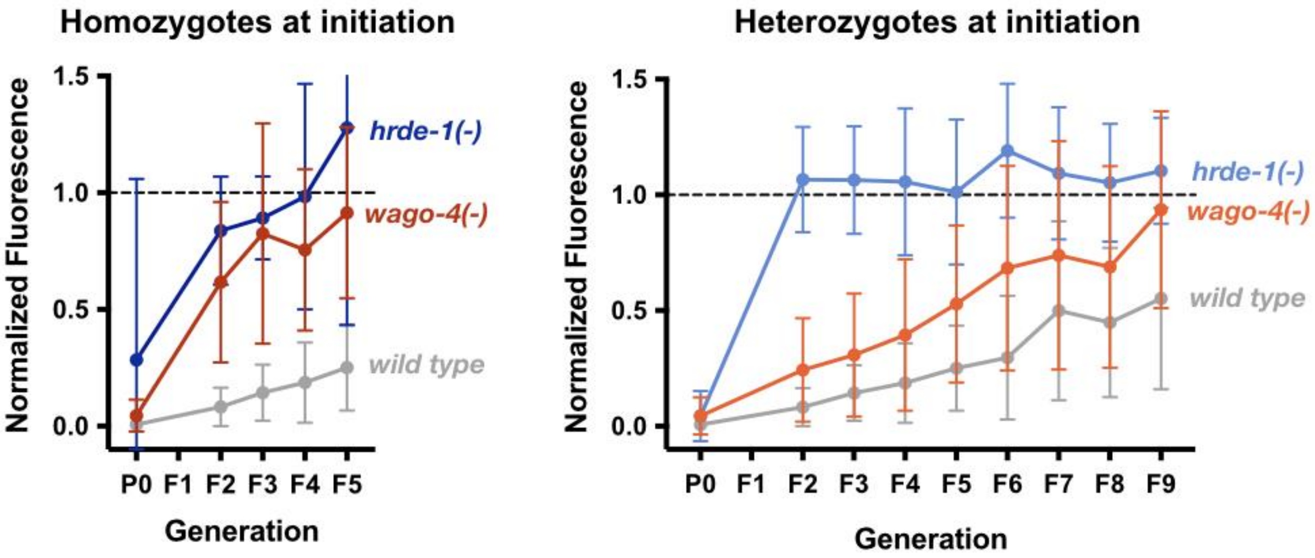


**Figure S4. *wago-4* mutants can inherit RNAi if functional *wago-4* is present at the initiation stage.**

Homozygote (left) and heterozygote (right) worms mutated in the argonautes *hrde-1* (blue) and *wago-4* (red) were exposed to *gfp* dsRNA for RNAi initiation, and GFP fluorescence (y-axis) was measure over generations (x-axis). The heterozygote worms (right) were heterozygote only at the P0 generation, in the next generations we measured GFP fluorescence in the homozygote mutant progeny. Fluorescence in each group was normalized to the mean fluorescence value of the corresponding isogenic control worms originally exposed to Empty vector. Shown are mean ± SD (30~120 worms per group) from two independent experiments combined together. All groups were tested side by side, therefore the same data of the wild-type control group appears on both panels.

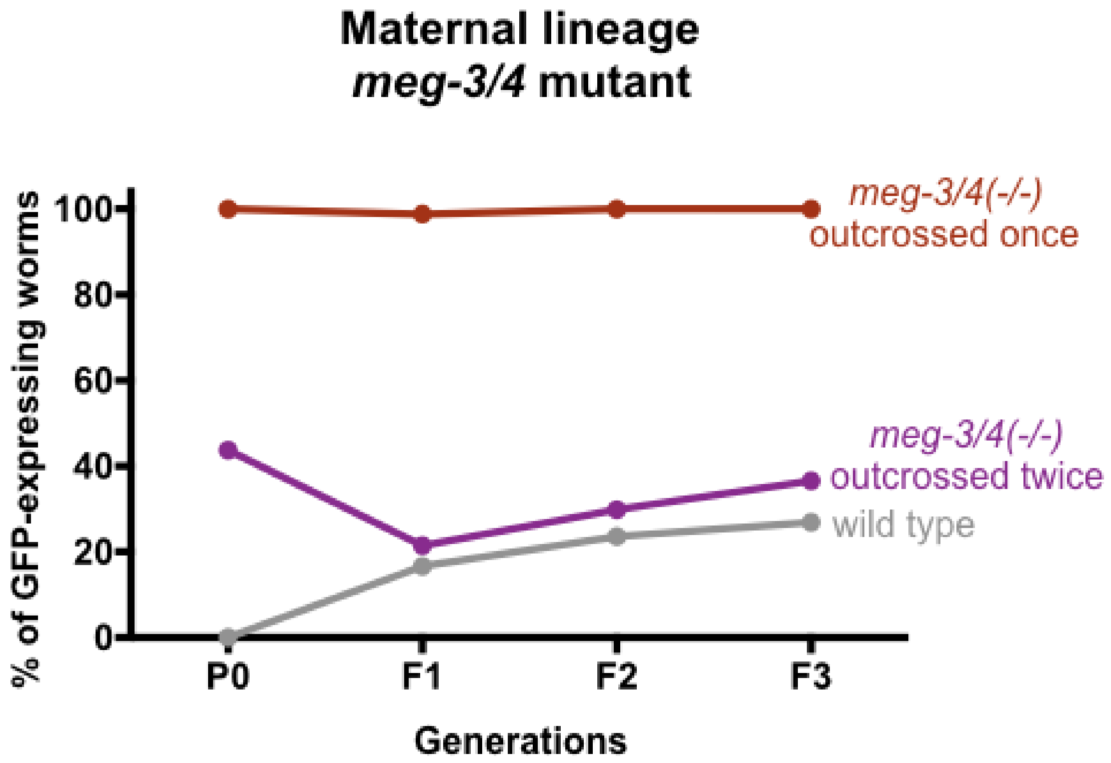


**Figure S5. Repetitive crossing of *meg-3/4* mutants to wild type males leads to improvement in the capacity of the mutants to generate an RNAi response.**

Worms with the indicated genotype were exposed to *gfp* dsRNA to generate RNAi two generations after homozygosity. The proportion of GFP-expressing worms (y-axis) was measured over generations (x-axis). *meg-3/4(-/-)* were outcrossed twice (purple, three generations passed between each outcross), and tested for RNAi inheritance. The experiment was performed side by side with a regular RNAi experiment after reciprocal crossing.

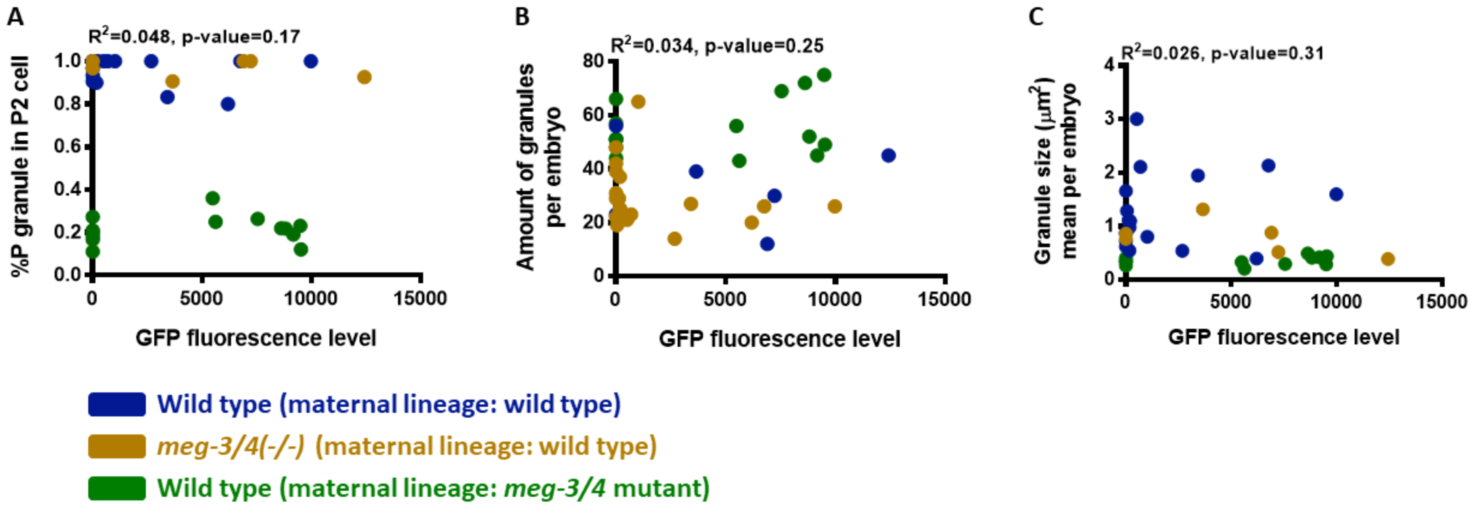


**Figure S6. The level of RNAi silencing does not correlate with P granule morphology.**

Worms of the indicated genotype and lineage were exposed to *gfp* dsRNA. Embryos exposed to RNAi were analyzed by microscopy. P granule characteristics (y-axis) are plotted against the GFP fluorescence levels (x-axis). Each dot represents one analyzed embryo. Pearson correlation scores and their P values are indicated above the panels.
